# Functional multispectral optoacoustic tomography imaging of hepatic steatosis development in mice

**DOI:** 10.1101/2020.12.17.419531

**Authors:** Shan Huang, Andreas Blutke, Annette Feuchtinger, Uwe Klemm, Robby Zachariah Tom, Susanna Hofmann, Andre C. Stiel, Vasilis Ntziachristos

## Abstract

The increasing worldwide prevalence of obesity, fatty liver diseases, as well as the emerging understanding of the importance of lipids in multi-faceted aspects of various other diseases is generating significant interest in lipid research. Lipid visualization in particular can play a critical role in understanding functional relations in lipid metabolism. In this study, we investigate the potential of multispectral optoacoustic tomography (MSOT) as a novel modality to non-invasively visualize lipids in laboratory mice. Using an obesity-induced non-alcoholic fatty liver disease (NAFLD) mouse model, we examined whether MSOT could detect and differentiate different grades of hepatic steatosis and monitor the accumulation of lipids in the liver quantitatively over time, without the use of contrast agents, *i.e.* in label free mode. Moreover, we demonstrate the efficacy of using the real-time clearance kinetics of indocyanine green (ICG) in the liver as a biomarker to evaluate the organs’ function and assess the severity of NAFLD. This study demonstrates MSOT as an efficient imaging tool for lipid visualization in preclinical studies, particularly for the assessment of NAFLD.

## Introduction

Non-alcoholic fatty liver disease (NAFLD) is characterized by excessive accumulation of lipids in the liver (1). Due to the increasing prevalence of obesity worldwide, NAFLD is rapidly becoming one of the leading causes of severe liver disease (2). Currently, the assessment of NAFLD severity relies on histopathology, which is not capable of monitoring the pathological and functional changes during therapeutic intervention without liver biopsies, limiting its use in pre-clinical studies. Thus, there is a need for a sensitive, quantitative, and non-invasive tool for NAFLD assessment in both preclinical and clinical settings, in order to enable longitudinally studies of potential therapies for this disease.

Ultrasonography (USG) is widely used as screening method for steatosis and fibrosis (3). However, it is not sensitive enough to detect mild steatosis. As of late, the use of magnetic resonance imaging to estimate the proton density fat fraction (MRI-PDFF) has been shown to more accurately and sensitively detect grades of steatosis in NAFLD than USG (4). Moreover, MRI-PDFF is responsive to changes in liver fat over time (5). Despite its value in clinical trials and experimental studies, MRI-PDFF application in the clinical setting is limited due to its high cost (6).

Currently, liver function assessment relies on blood tests for classic biomarkers that are poorly correlated with NAFLD development, such as aspartate aminotransferase (AST) and alanine aminotransferase (ALT) (7). Beyond that, the indocyanine green (ICG) clearance test is considered a valuable method for dynamic assessment of liver function (8). ICG is a water-soluble fluorescent dye that is exclusively cleared from the blood via the liver. The clearance of ICG through the hepatobiliary tract depends on liver blood flow and hepatic and biliary function (9). Thus, its clearance from the blood stream serves as a biomarker of liver function (10), making the ICG clearance test potentially valuable for NAFLD monitoring (11) (12). However, the readout of clinical tests of ICG clearance from the blood stream cannot reflect the full process of ICG clearance in the liver. Using a rabbit model, Seifalian et al. demonstrated that the ICG uptake rate correlated significantly with blood flow and microcirculation, while the excretion rate correlated with liver enzymes, which reflected the function of hepatocytes (11). Since NAFLD starts with lipid accumulation in hepatocytes, the early changes in liver function caused by NAFLD should affect the excretion of ICG from the hepatocytes to the bile ducts. Therefore, blood ICG clearance, which reflects mainly the uptake of ICG by the liver rather than the excretion of ICG from the hepatocytes, might not be sensitive enough for the detection of early changes in liver function during NAFLD development. This insensitivity may also underlie the skepticism in some studies, as to whether the ICG clearance rate correlates with the severity of steatosis (12) (13) (14).

Multi-spectral optoacoustic tomography (MSOT) is a novel video-rate imaging modality that can directly visualize lipids based on their characteristic absorption of light at approximately 930 nm, a wavelength that falls in the near-infrared (NIR) spectral range. The technique detects ultrasound waves that are generated by thermo-elastic expansion of tissue upon absorption of light by different biomolecules, such as lipids. The technique combines the molecular sensitivity of light with the resolution of ultrasound, yielding a potent technology for the study of lipids. Due to the low attenuation of light in the NIR compared to the visible or infrared spectral ranges, operation at 930 nm allows light penetration through several centimetres of tissue, making MSOT highly effective for *in vivo* imaging (15) (16). Moreover, MSOT is sensitive to ICG at the dye’s peak absorption at ~800 nm, potentially allowing both lipids and ICG distribution and dynamics to be monitored (17) (18). Some studies have applied optoacoustic (OA, also known as photoacoustic) methods to investigate steatosis *in vivo.* In contrast to MSOT, which uses the full spectral information to achieve molecular sensitivity, these studies used single wavelength information to detect steatosis based on the lipid absorption signature (19) (20). However, none of these studies has compared their OA-based approach with the gold standard histopathology for steatosis diagnosis, limiting their value for clinical translation.

We hypothesized that MSOT could be used to non-invasively evaluate steatosis by directly imaging and quantifying changes in hepatic lipid content of mice *in vivo.* As a first step, we used MSOT to characterize lipid content, as well as water, oxy- and deoxy-haemoglobin (HbO_2_ and Hb) content of various tissues *in vivo*, including liver, kidney, brown adipose tissue (BAT), white adipose tissue (WAT), aorta, and sulzer vein (SV). We then assessed the ability of MSOT to detect excessive lipid in the liver of a NAFLD mouse model *in vivo* with quantitative readouts, and compared our data to semi-quantitative histopathological grading, the current gold standard for evaluation of hepatic steatosis. To demonstrate the performance of our MSOT-based methods in predicting grades of hepatic steatosis, we analysed the receiver operating characteristic (ROC) curve. We further monitored the progression of steatosis in mice over time using our MSOT imaging approach. Lastly, we tested the feasibility of assessing hepatic function by detecting ICG clearance kinetics based on OA detection of ICG directly in the liver. Our results introduce MSOT as a novel imaging tool for NAFLD assessment in pre-clinical studies, which holds potential for monitoring of NAFLD in clinical settings.

## Materials and Methods

### Animals

Animals were kept at 24±1°C and on a 12:12-h light-dark cycle with free access to food and water. To induce obesity and NAFLD, 8-10 week old male C57BL/6BrdCrHsd-Tyrc mice (Janvier Labs, France; The Jackson Laboratory, United States) were fed with HFD comprising 58% kcal from fat (D12331; Research Diet, New Brunswick, NJ, USA) for up to 6 months. Control mice from the same strain were fed with standard rodent diet (Altromin 1314, Altromin Spezialfutter GmbH & Co, Germany). For the mice used in the ICG experiment, 2.5μg/g of ICG (ICG-Paulsion von Verdye 25mg, Diagnostic Green, Germany and USA) in 100μl saline was injected i.v. through a catheter. All MSOT imaging experiments were carried out at 34°C. The animal studies were approved and conducted in accordance to the Animal Ethics Committee of the government of Upper Bavaria, Germany.

### Multi-spectral Optoacoustic Tomography

MSOT measurement were conducted with a 256-channel real-time imaging MSOT scanner (inVision 256, iThera Medical, Germany) described before (15) (21). For *in vivo* measurement, mice were anesthetized by continuous inhalation of 2% isoflurane throughout imaging. Then the mice were placed onto a thin, clear, polyethylene membrane and positioned in the water bath in a sample holder as described earlier (22).

For *ex vivo* measurement, tissues were isolated from the mice right after euthanasia. Samples were placed into 1 mL syringes filled with PBS and held in position by a custom designed holder in MSOT. Imaging was performed at 27 wavelengths in the range of 700 nm-960 nm in step of 10 nm.

### Histopathology

Liver tissue specimen were sampled according to established organ sampling and trimming guidelines for rodent animal models (23). The samples were fixed in neutrally-buffered 4% formaldehyde solution for 24 hours and subsequently routinely embedded in paraffin. 3 μm thick sections were stained with haematoxylin and eosin (HE), using a HistoCore SPECTRA ST automated slide stainer (Leica, Germany) with prefabricated staining reagents (HistoCore Spectra H&E Stain System S1, Leica, Germany), according to the manufacturer’s instructions. Histopathological examination was performed by a pathologist in a blinded fashion (*i.e.*, without knowledge of the treatment-group affiliations of the examined slides). Hepatic steatosis was graded semiquantitatively (24) according to the proportion of liver tissue in the sections affected by the presence of fat vacuoles in hepatocyte profiles (grade 0: < 5%; grade 1: 5-33%; grade 2: 33-66%; grade 3: > 66%).

### Oil-red-O staining

Livers were isolated right after euthanasia. Tissues were embedded in O.C.T. (TissueTeck; Sakura Finetek, USA) and frozen to −50°C before cryosliced with a step of 10 μm at −20 °C using a modified cryotome (CM 1950; 839 Leica Microsystems, Germany). The slices were air-dried for 2 hours before staining. The slices were first fixed in neutrally-buffered 4% formaldehyde solution, briefly washed with running tap water for 1 - 10 minutes and rinsed with 60% isopropanol. Then the slices were stained with Oil-Red-O working solution, freshly prepared according to the manufacture’s manual (O0625-25G, Signa-Aldrich, Merck KGaA, Darmstadt, Germany) for 15 minutes followed by rinse with 60% isopropanol. Then the nuclei were lightly counter-stained with alum haematoxylin. The slices were finally rinsed with distilled water before mounting of cover slips with aqueous mountant (P36971, ThermoFisher SCIENTIFIC, Waltham, Massachusetts, United States).

### Data analysis

MSOT data were analysed by ViewMSOT software (v3.8, iThera Medical, Munich, Germany). MSOT images were reconstructed using the model linear method. For unmixing of Hb, HbO_2_, Lipid, H_2_O, and ICG, a linear regression method was used. For detection of lipid, difference unmixing was also applied. The data analysis additionally consisted of the calculation of the total blood volume (TBV = HbO_2_+ Hb, displayed as arbitrary units) and the oxygen saturation (sO_2_= HbO_2_ / TBV, displayed as percentage). Each unmixing data point for statistics was averaged from three ROIs in the same subject. To normalize the ICG intensity data, we subtracted the background signal of the ROI in the liver from the raw unmixing data before ICG administration, and then normalized the data to the peak value.

### *Ex vivo* quantification of ICG tracer

Dissected liver samples were snap frozen in liquid nitrogen, cryosectioned at 12 μm (CM1950, Leica Microsystems, Wetzlar, Germany), counterstained with H33342 dye (H33342; Sigma Aldrich, Merck KGaA, Darmstadt, Germany) and mounted with Vectashield (H-1000, Biozol Eching, Germany). Stained tissue sections were scanned with an AxioScan.Z1 digital slide scanner (Zeiss, Jena, Germany) equipped with a 20x magnification objective. ICG based signals were detected with a filter set FT 762 - BP785/38 and H33342 with a filter set FT 405 - BP 425/30. Images of the entire tissue sections were acquired and evaluated using the commercially available image analysis software Definiens Developer XD 2 (Definiens AG, Munich, Germany). The calculated parameter was the mean ICG fluorescence intensity of the whole liver tissue section.

### Statistics

Data were analyzed using GraphPad Prism (v. 8.4.2; GraphPad Software, La Jolla, CA) except for Receiver Operating Characteristics (ROC) analysis, which was performed using easyROC (v. 1.3.1; easyROC, www.biosoft.hacettepe.edu.tr/easyROC). All data presented as mean ± SEM unless elsewhere stated. Group size (n) is indicated for each experiment in figure legends. Student’s t-test was used for comparisons of 2 independent groups. One-way ANOVA followed by Tukey’s post hoc test was used for comparing >2 independent groups. The *z* test was used for comparisons of AUROC between 2 groups (25). In ROC analysis, optimal cutoff is determined by Youden’s index. A p-value of < 0.05 was considered as statistically significant. Significant digit: *P < 0.05, **P < 0.01, *** P < 0.001

## Results

### MSOT imaging of liver, kidney, adipose tissues, and blood vessels in non-obese mice

First, we examined the ability of MSOT to differentiate various key tissues based on the spectral signatures of HbO_2_, Hb, lipid, and water in healthy mice, which is the prerequisite for tissue identification and eventual pathophysiological observation. Hb and HbO_2_ can be used to assess metabolic metrics, such as total blood volume (TBV) and oxygenation of the tissue (sO_2_). The right kidney serves as a negative control for the quantification of fat in the liver, as it is located close to the liver and does not accumulate large amounts of fat during obesity-related development of hepatic steatosis. Interscapular brown adipose tissue (iBAT) and retroperitoneal white adipose tissue (rpWAT), were positive controls for the lipid detection. The aorta and SV, were also examined, mainly for tissue oxygenation analysis due to their pure blood content and natural difference in sO_2_. Inspecting the anatomical and spectral features of certain tissues in single wavelength OA images at 800 nm and corresponding images with unmixing data of tissue contents (Figure 1A), we determined the region of interest (ROI) of liver, kidney, iBAT, rpWAT, aorta, and SV in mice. Then we checked the OA features of the tissues by inspecting their absorption spectra (Figure S1A-C) in comparison with individual endogenous absorbers (Figure S1D). To enhance the visualization of spectral features, we normalized all tissue spectra to the highest OA signal acquired in the 700-960 nm range (Figure 1B). Since the *in vivo* spectra of a deep tissue can be affected by the surrounding tissues due to light attenuation, we compared *in vivo* spectra of liver, kidney, iBAT and rpWAT with their *ex vivo* spectra from the same animal and found no remarkable difference between the *in vivo* and *ex vivo* spectra (Figure 1C).

**Fig.1.**
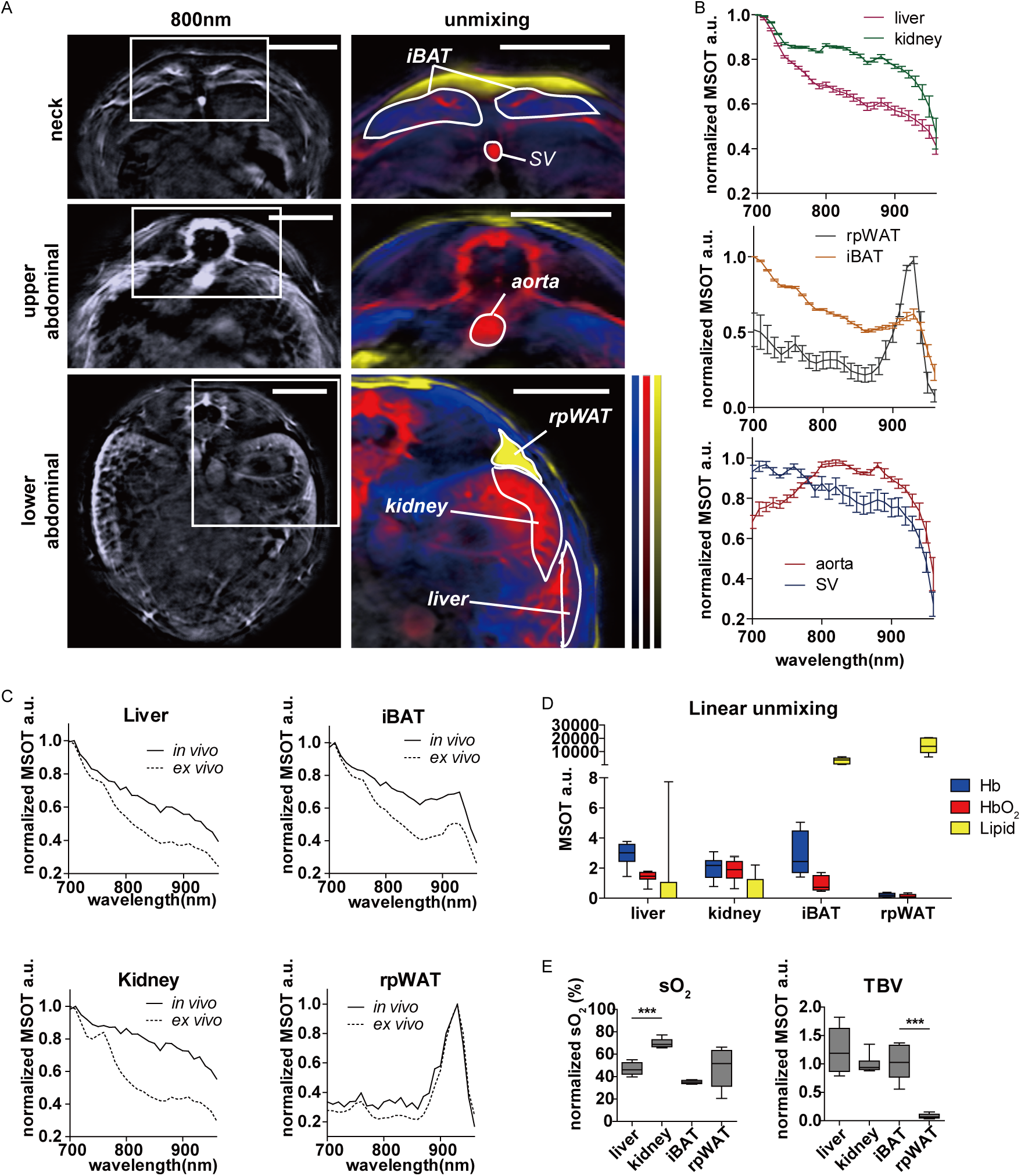
MSOT imaging of liver, kidney, interscapular brown adipose tissue (iBAT), retroperitoneal white adipose tissue (rpWAT), aorta, and sulzer vein (SV) *in vivo* and *ex vivo.* A, Reconstructed MSOT image (800 nm) without and with linear unmixing data in neck, upper and lower abdominal area showing of liver, kidney, iBAT, rpWAT, aorta, and SV ROIs. Unmixing result: blue for Hb, red for HbO_2_, yellow for lipid. The color bar shows the color coding of MSOT a.u. from minimum to maximum (bottom to top) Scale bar: 4mm. B. Normalized spectra of liver, kidney, iBAT, rpWAT, aorta, and SV. n = 8. C. Normalized spectra of liver, kidney, iBAT, and rpWAT *in vivo* and *ex vivo.*D-E, Box and whiskers (whiskers: min to max) of linear unmixing results from liver, kidney, iBAT, and rpWAT. n = 8.

In the following quantitative analysis, we normalized the tissue TBV and sO_2_ to the respective aorta readouts of each mouse. As expected, liver and kidney were detected as blood-rich tissues (Figure 1D,E). Due to the double vascularization system in the liver, which contains 80% venous blood from the portal vein (26), the sO_2_ in liver is significantly lower than that in kidney (P < 0.0001). In comparison to rpWAT, iBAT has higher blood content (p < 0.0001) (Figure 1E), which reflects its high vascular density (27).

These results demonstrated that MSOT is suitable to analyse the contents and the metabolic status of key tissues, including liver, kidney, iBAT, and rpWAT *in vivo.*

### MSOT imaging of steatosis in mice with diet-induced obesity

After confirming the ability of MSOT to differentiate the key tissues in healthy animals, we assessed whether pathological changes related to fat accumulation alter the spectral signature of the tissues. We chose the diet-induced obesity mouse model which develops steatosis along with obesity (28). The mice fed with high fat diet (HFD) had enlarged fat depots and significantly higher body weight (P=0.005) (Figure 2A, B). While we saw no significant change in the concentration of Hb and HbO_2_ in any of the tissues analysed, the lipid content significantly increased in the liver (P=0.004) and iBAT (P=0.007) (Figure 2C). This increase indicates a synergistic malfunction in lipid metabolism in these tissues, consistent with similar findings from recent studies (29) (30). The metabolic status of the tissues revealed by sO_2_ and TBV was not altered in the obese mice (Figure 2D). These results suggest that MSOT, without labels, is able to assess pathological changes associated with steatosis.

**Fig.2.**
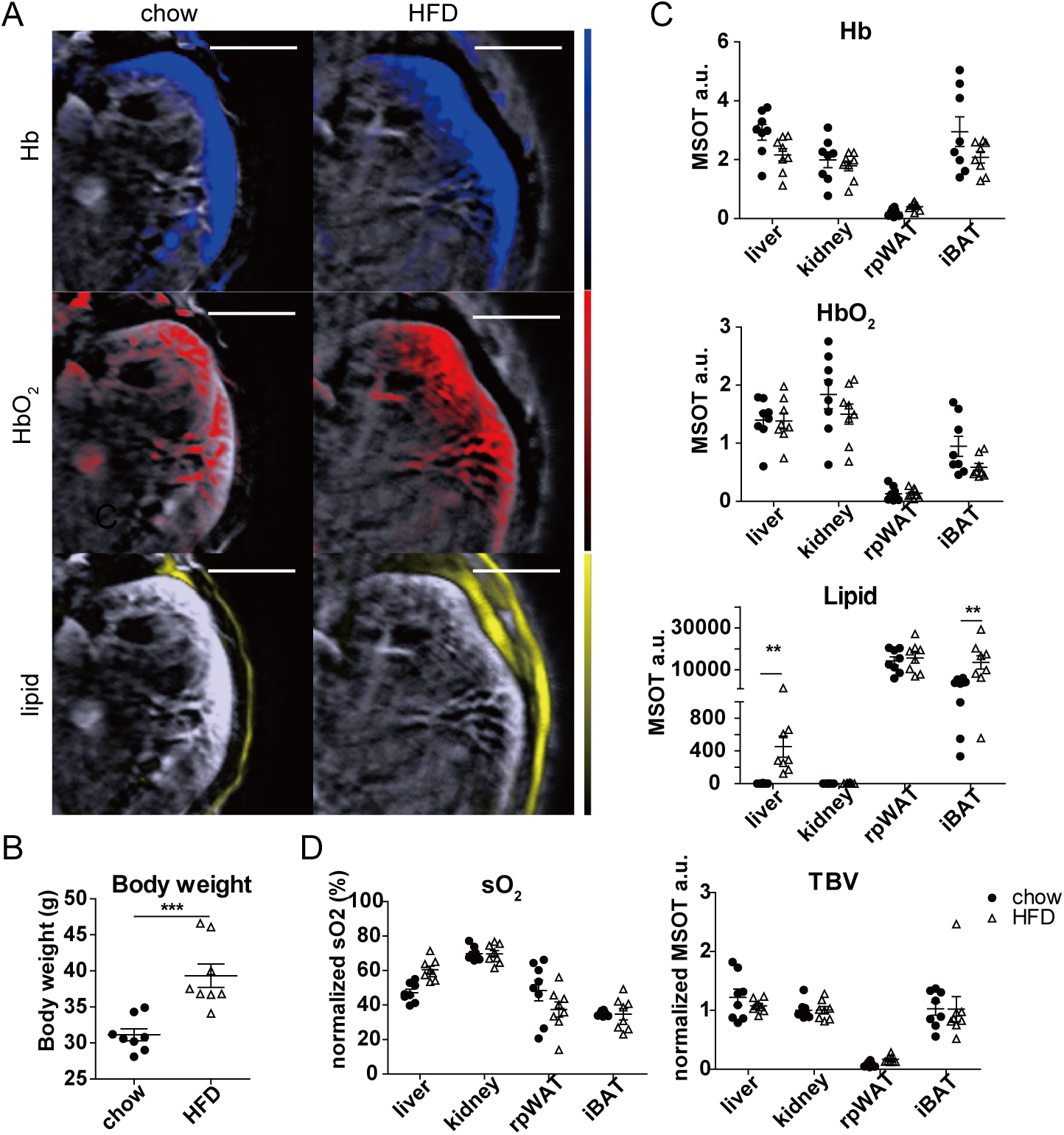
Comparison of oxygenation and lipid content of target tissues between healthy and obese mice using MSOT imaging. A, Reconstructed MSOT image (800 nm) with linear unmixing data of Hb, HbO_2_, and lipid from chow and HFD-fed mice. The color bar shows the color coding of MSOT a.u. from minimum to maximum (bottom to top). Scale bar: 4 mm. B, Body weight of chow- and HFD-fed mice. n = 8. C, Unmixing result of Hb, HbO_2_ and lipid from liver, kidney, iBAT, and rpWAT, n = 8. D, Tissue oxygenation (sO_2_) and total blood volume (TBV) results from liver, kidney, iBAT, and rpWAT, n = 8.

### Specific and sensitive detection and quantification of lipid in liver by MSOT

Here we verified the specificity and sensitivity of lipid detection in normal and steatotic liver by MSOT. Separate from the linear unmixing method, we introduced another method for label-free lipid detection based on the absorption difference between 700 nm and 930 nm. To demonstrate the two methods, we first imaged the livers of one lean mouse and one obese mouse with extreme steatosis (confirmed by Oil-red-O lipid staining) both *in vivo* and *ex vivo* (Figure 3A, B). The tissue spectra of the steatotic liver showed strong signals at 930 nm (indicative of lipid) when compared to the control, both *in vivo* and *ex vivo* (Figure 3C). In comparison, the spectra of the kidneys *in vivo* in the same subjects were nearly identical, as expected (Figure 3C). We used the intense signal 930 nm in the steatotic liver to derive a simpler numeric readout for lipid detection than that of linear unmixing. This was accomplished by subtracting the intensity at 930 nm from that at 700 nm (primarily blood absorption). This value was lower in the steatotic liver than in the normal liver *in vivo* (Figure 3B, bottom) and *ex vivo* (Figure 3D). For the quantification, we normalized the difference value to the signal at 800 nm to minimize the individual variance. Furthermore, we quantified the lipid content from 6 healthy and 9 NAFLD mice with severe steatosis. The average lipid contents calculated by linear unmixing were 1.29 a.u. and 674.6 a.u. (P = 0.0358) in the control and steatotic livers, respectively, while the corresponding readouts of (700 nm-930 nm)/800 nm was 0.772 and 0.396 (P = 0.0008) (Figure 3E). Using both linear and difference unmixing, the readouts from the adjacent right kidney were not affected by the hepatic steatosis (Figure 3E). For later performance analysis, we divided (700 nm - 930 nm)/800 nm readout from kidney by that from liver to further eliminate the individual variance while achieving positive relevance with steatosis. The final readout is named as difference index.

**Fig.3.**
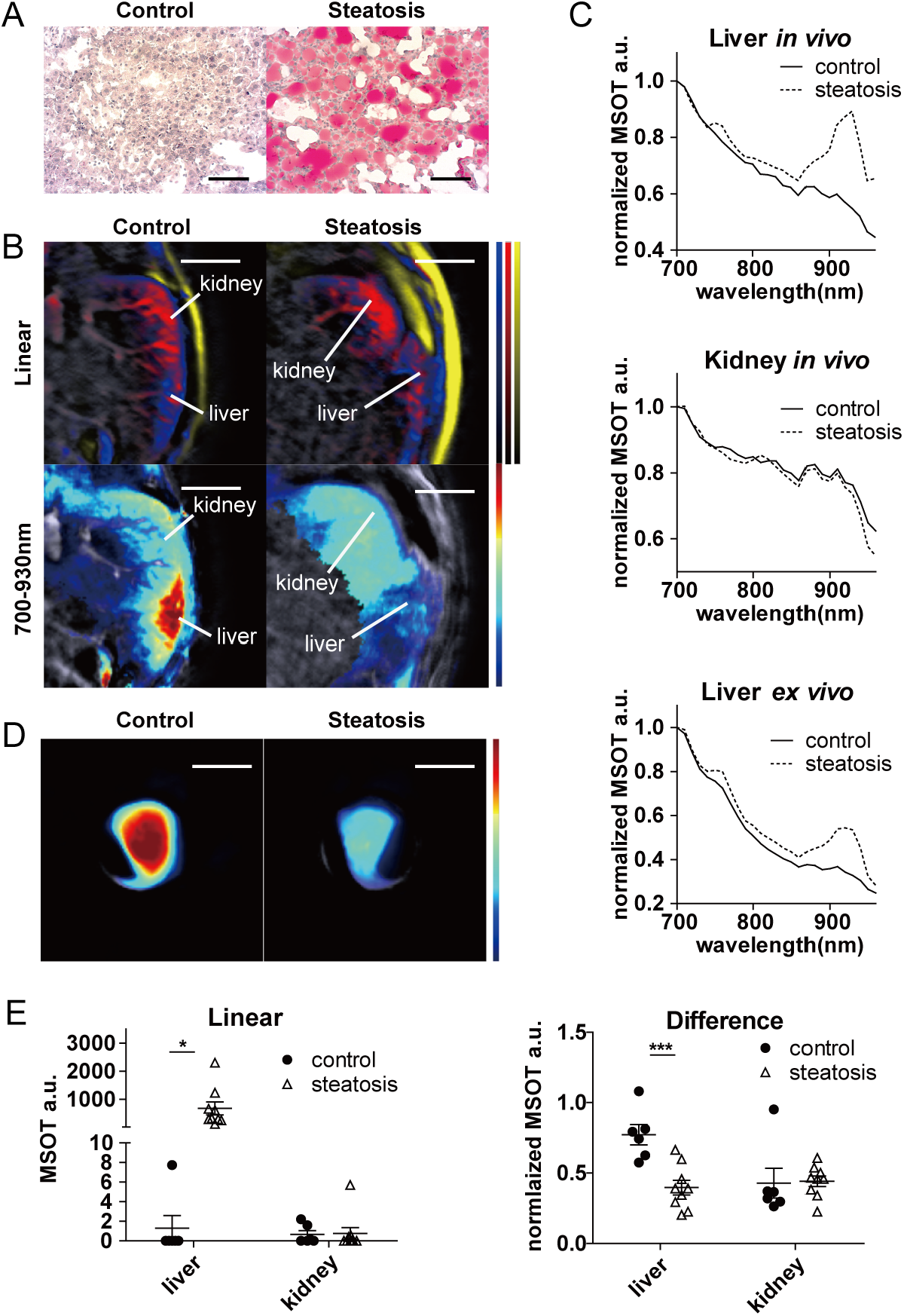
MSOT detection of lipid content in healthy and steatotic livers *in vivo* and *ex vivo.* A, Oil-red-O staining of control (healthy) and steatotic liver. Scale bar: 100 μm. B, Reconstructed MSOT image (800 nm) with linear unmixing and 700 nm – 930 nm difference data of lipid in lower abdominal section. Unmixing result: blue for Hb, red for HbO_2_, yellow for lipid, jet for 700 nm – 930 nm difference. The color bar shows the color coding of MSOT a.u. from minimum to maximum (bottom to top) Scale bar: 4 mm. C, Normalized spectra of livers and kidneys from control (healthy) and subject with hepatic steatosis. D, Reconstructed MSOT image (800 nm) with linear and 700 nm – 930 nm difference unmixing data of lipid in control (healthy) and steatotic liver *ex vivo.* The color bar shows the color coding of MSOT a.u. from minimum to maximum (bottom to top) Scale bar: 4 mm. E. Linear and difference data of lipid from control and steatotic (grade 3) livers. Control: n = 6, Steatosis n= 9. Data in panel A-D are from the same subjects.

Furthermore, we examined the performance of both methods in lipid quantification and compared the results with histopathological grading. Representative examples of the full range of grades of steatosis are shown in Figure 4A. By linear unmixing of the spectra measured by MSOT, the difference between grades 3 and grade 0 was highly significant (P < 0.0001) (Figure 4B). Moreover, linear unmixing is also able to distinguish grade 1 or 2 and grade 3 steatosis (grade 1 vs 3: P = 0.0002; grade 2 vs 3: P=0.004) (Figure 4B). Livers with all grades of steatosis had significantly higher difference index values than the normal liver (grade 0 vs 1: P = 0.0007; grade 0 vs 2: P = 0.0009; grade 0 vs 3: P < 0.0001) (Figure 4C).

**Fig.4.**
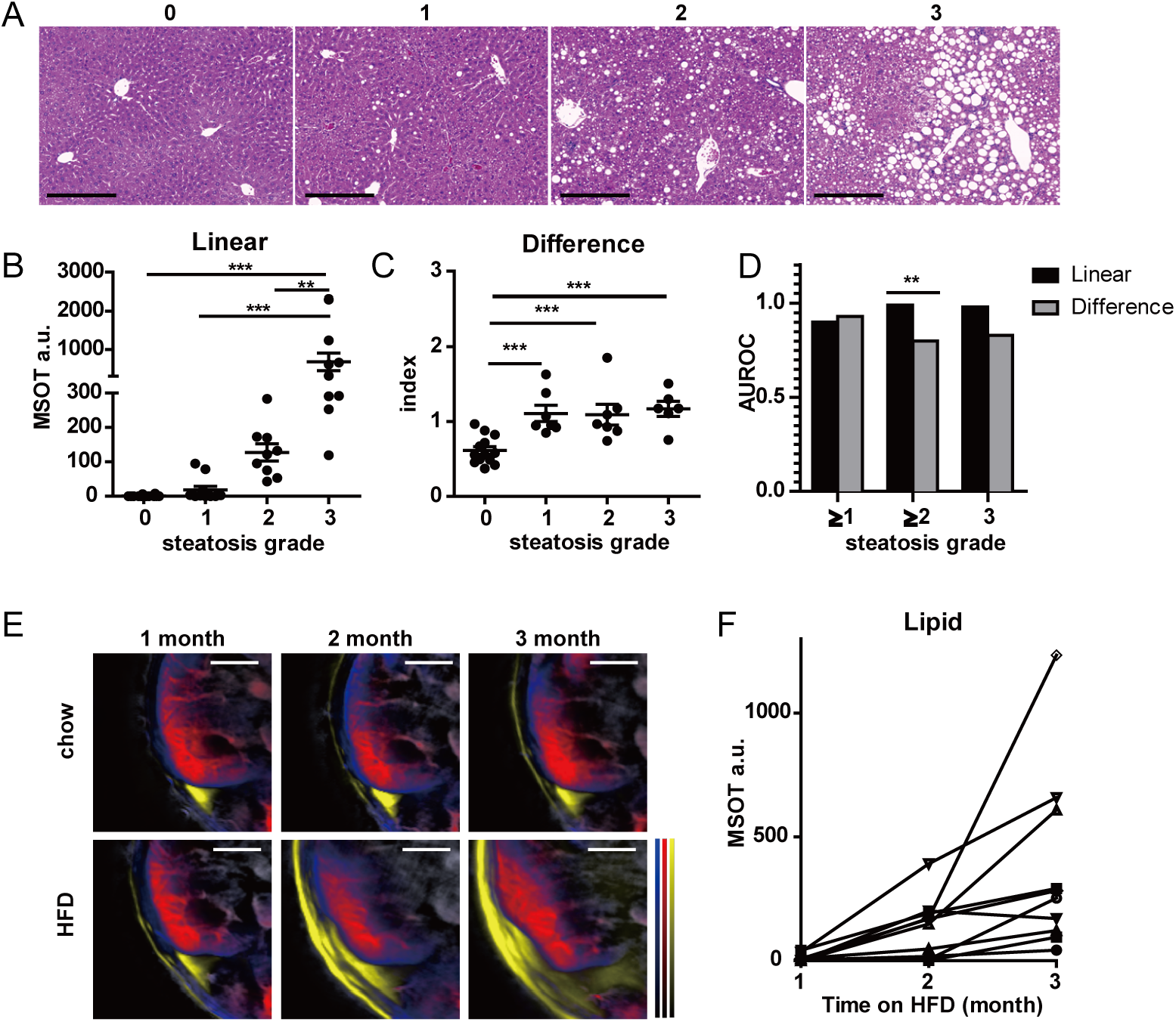
Quantitative assessment of steatosis by MSOT in comparison to histological grading. A, HE staining of livers with grade 0 – 3 steatosis. Scale bar: 100 μm. B, Quantification of linear unmixing of lipid in liver. Grade 0: n = 14; grade 1: n = 11; grade 2 & 3: n = 9. C, Quantification of difference index in liver. Same subjects as in B. D, Diagnostic accuracy of linear and difference unmixing in diagnosing dichotomized grades of steatosis. E. Reconstructed MSOT image (800 nm) with linear unmixing data of Hb, HbO_2_, and lipid from chow and HFD-fed mice. Unmixing result: blue for Hb, red for HbO_2_, yellow for lipid. The color bar shows the color coding of MSOT a.u. from minimum to maximum (bottom to top). Scale bar: 4 mm. F. Longitudinal track of lipid in mouse livers. Each line represents data from one animal. All animals have grade 3 steatosis at the endpoint confirmed by histology.

To investigate the prediction accuracy of MSOT readouts for steatosis in preclinical setting, we calculated the area under the receiver operating characteristic (AUROC) and the potential cutoff value for assessing different grade of steatosis (Figure 4D, Table 1). The ROC analysis results suggested that both linear and difference unmixing can predict grades of steatosis with excellent (AUROC>0.8) to outstanding (AUROC>0.9) discriminatory ability (31). Direct comparison showed that linear unmixing was more accurate than difference unmixing at predicting grade 2-3 steatosis (P=0.008, Figure 4D).

**Table 1.** Prediction Accuracy of linear unmixing and difference unmixing in Detecting Each Grade of Steatosis. Se, sensitivity; Sp, specificity. NPV, negative predictive value; PPV, positive predictive value.

| Cross-sectional<br>steatosis grade<br>classification (n=43) | linear unmixing |  |  |  |  |  | Lipid index ((700-930)/800nm kidney/liver) |  |  |  |  |  | vs Linear<br>unmixing<br>P value |
| --- | --- | --- | --- | --- | --- | --- | --- | --- | --- | --- | --- | --- | --- |
|  | cutoff | AUROC<br>(95%CI) | Se | Sp | PPV | NPV | cutoff | AUROC<br>(95%CI) | Se | Sp | PPV | NPV |  |
| Grade 1-3 (n=29) vs<br>Grade 0 (n=14) | 17.3 | 0.90<br>(0.81-0.98) | 72% | 100% | 100% | 64% | 0.74 | 0.93<br>(0.86-1.02) | 100% | 79% | 91% | 100% | 0.48 |
| Grade 2-3 (n=18) vs<br>Grade 0-1 (n=25) | 42.9 | 0.99<br>(0.96-1.01) | 100% | 92% | 90% | 100% | 0.97 | 0.80<br>(0.67-0.94) | 67% | 84% | 75% | 78% | 0.008 |
| Grade 3 (n=9) vs<br>Grade 0-2 (n=34) | 252.9 | 0.98<br>(0.94-1.02) | 89% | 97% | 89% | 97% | 1.01 | 0.83<br>(0.68-0.98) | 89% | 82% | 57% | 97% | 0.06 |

We further tested the possibility of tracking the progression of steatosis *in vivo.* HFD-fed mice gradually accumulated excessive lipids in fat depots (Figure 4E). Despite individual differences in actual values, the measured hepatic lipid levels all exhibited an upward trend during the 3 months of HFD feeding, indicating a gradual accumulation of lipid in liver over time (Figure 4F). These results suggested the feasibility of monitoring steatosis progression by MSOT in a preclinical setting.

### Functional OA imaging of liver with ICG

Here we tested our hypothesis that hepatic ICG clearance could serve as a biomarker for liver function in the assessment of NAFLD in mice. After administration of ICG, we traced its levels based on linear unmixing in healthy and steatotic livers for 2 hours, before which the normal liver clearance of ICG was completed (Figure 5A, S2A, Movie S1). The liver spectra showed peak absorption at 800nm in both control and steatotic liver, indicating existence of ICG in the tissue after injection. The 800nm peak disappeared in the control liver 2 hours after ICG injection, while in steatotic liver, the peak remained visible throughout the observation (Figure 5B, S2B). We further monitored the kinetics of ICG signal for 1 hour after ICG administration in 3 normal and 3 NAFLD mice with grade 3 steatosis, confirmed by histology and MSOT imaging (P = 0.0003) (Figure 5C). The steatotic livers showed slower elimination of ICG than the normal liver (Figure 5D), indicating an impaired excretion function. Consistently, there was more residual ICG signal in the steatotic liver than in the normal liver under fluorescent microscopy *ex vivo.* (Figure 5E, S3).

**Fig.5.**
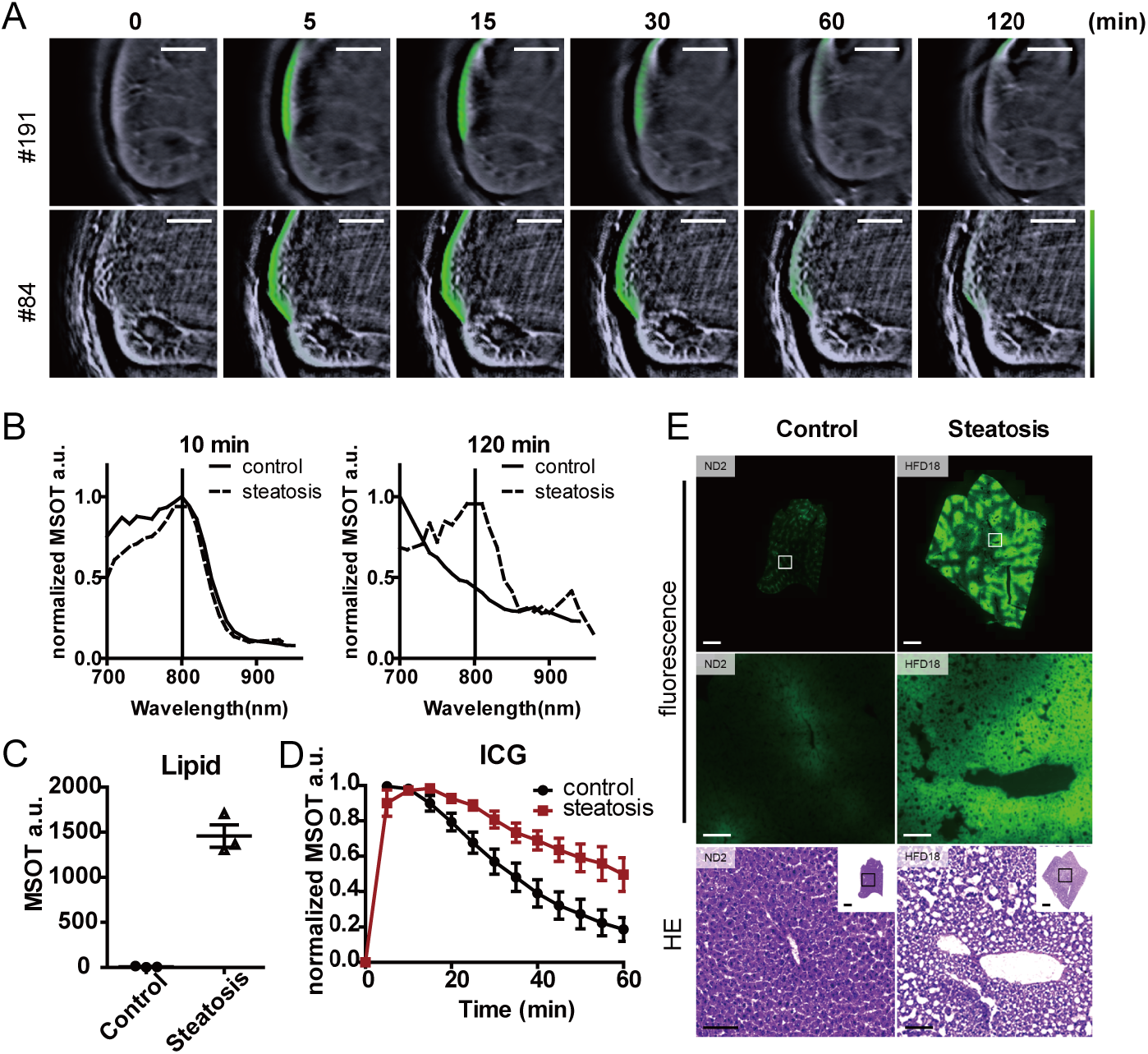
MSOT imaging of ICG clearance in mouse livers. A, Reconstructed MSOT image (800 nm) with linear unmixing data of ICG from mice with and without steatosis. Scale bar: 4 mm. B. Normalized spectra of control and steatotic liver 10 and 120 min after ICG injection. C, Quantification of linear unmixing of lipid in livers with and without steatosis. D, Longitudinal monitoring of ICG intensity in control and steatotic livers (time interval: 5 minutes; total length: 60 minutes). n = 3. For each curve, data is normalized to the highest intensity acquired. E, Representative fluorescence microscopy and HE staining images of control and steatotic liver. Error bar: 100 μm.

## Discussion

Few studies use label-free OA imaging methods to investigate the biology and pathology of the liver (19) (20), most likely due to the high absorption of light by blood and the depth. Herein we demonstrate that by exploiting its multispectral nature, MSOT imaging is able to detect and separate all grades of steatosis with quantitative readouts, supporting its value for disease progression and treatment efficacy monitoring in a pre-clinical setting. Compared to linear unmixing, the difference unmixing showed similar sensitivity as linear unmixing towards mild steatosis, despite its use of only three wavelengths (700, 800 and 930 nm). For a conceivable clinical translation, this approach would be faster, more straight-forward to analyse, and requiring less cost intensive laser sources than linear unmixing, making it suitable for screening and early detection of steatosis.

In this work, we have also introduced hepatic ICG clearance as a biomarker for NAFLD assessment in mice using time-lapse MSOT imaging. With the time-resolved data acquired by MSOT, we were able to differentiate the kinetics of ICG uptake and excretion in control and steatotic livers in mice. A similar method using tissue NIR spectroscopy has been introduced as a comprehensive liver function test, but its application was restrained by the invasive procedures including an open surgery (32). Our non-invasive method can overcome this limitation while providing equivalent data. Since ICG is a proven contrast agent for clinical use, it is feasible to test our MSOT-based ICG detect method in a clinical setting.

Despite this study aiming only at the analysis of steatosis in mice, the encouraging results in the use of MSOT in measuring spectral data in humans (15) suggest the possibility of a clinical translation of the method described in this work. In contrast to MRI, clinical MSOT is a portable modality, which could enable bed-side or outpatient monitoring of NAFLD development. However, this method needs to be further adapted for clinical translation. First, since we were using pre-clinical small-animal MSOT throughout this study, the method introduced here needs to be adjusted for the clinical version of MSOT (15). Second, the desired imaging depth in humans, which is commonly several centimetres, could present a challenge to preserve the sensitivity and the capability for quantification by our lipid detection method, especially in morbidly obese patients. However, intravascular imaging of lipids, which has been demonstrated in many studies for plaque detection (33) (34), might allow access to the periportal area of the liver through catheter based OA imaging. This would overcome the limitation of imaging depth and potentially lead to a more sensitive method for steatosis detection.

Clinical applications of optoacoustic imaging are rapidly emerging, with studies suggesting the potential of MSOT for diagnosing skin tumors (35), breast cancer (36), vascular malformations (37), systemic sclerosis (38), Crohn’s disease (39), and muscular dystrophy (40). Here we introduce MSOT as a new tool for NAFLD assessment in preclinical settings. Outcomes of future clinical trials may confirm the potential of MSOT as a sensitive, quantitative, and cost-effective imaging method for NAFLD assessment, which will ease the monitoring of the disease, as well as promote its therapeutic study for better healthcare outcomes.

## Author Contributions

S.H. conceived the study, designed and performed the experiments, analysed the data and wrote the manuscript. A.B. designed and performed the *ex vivo* experiments and analysed the data. A.F. designed and performed the *ex vivo* experiments and analysed the data. U.K. performed the *in vivo* experiments. R.Z.T performed the *in vivo* experiments. S.M.H conceived and oversaw the animal study and provided advice for physiological data interpretation and manuscript composition. A.C.S conceived the study and wrote the manuscript. V.N. conceived the study and wrote the manuscript.

## Acknowledgements

The authors would like to thank Sarah Glasl and Pia Anzenhofer for technical assistance, Doris Bengel for support in animal experiments and Robert J. Wilson for critical discussions on the manuscript.

## Competing Interests Statement

V.N. is a shareholder of iThera Medical.

## List of Abbreviations

AUROC: area under the receiver operating characteristic; Hb: deoxygenated blood; HbO_2:_ oxygenated blood; HFD: high fat diet; iBAT: interscapular brown adipose tissue; ICG: indocyanine green; MSOT: multispectral optoacoustic tomography; OA: optoacoustic; ROI: region of interest; rpWAT: retroperitoneal white adipose tissue; sO_2:_ tissue oxygenation; TBV: total blood volume

## Financial Support

The research leading to these results has received funding from the Deutsche Forschungsgemeinschaft (DFG), Germany [Gottfried Wilhelm Leibniz Prize 2013; NT 3/10-1], from the European Research Council (ERC) under the European Union’s Horizon 2020 research and innovation programme under grant agreement No 694968 (PREMSOT) and from the Helmholtz-Gemeinschaft Deutscher Forschungszentren (HGF)/ExNet project “Innovative Intelligent Imaging” (i3-Helmholtz).

