## Supplementary figure for "Functional multispectral optoacoustic tomography imaging of hepatic steatosis development in mice"

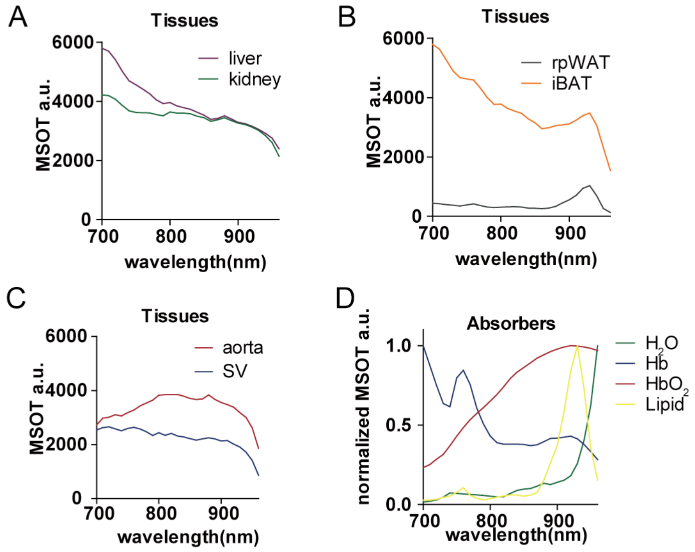


Fig.S1 MSOT spectra of liver, kidney, interscapular brown adipose tissue (iBAT), retroperitoneal white adipose tissue (rpWAT), aorta, and sulzer vein (SV) in vivo and main absorbers. A-C, Raw spectra of liver, kidney, iBAT, rpWAT aorta and SV in vivo. D, Normalized spectra of H2O, Hb, HbO2, and Lipid

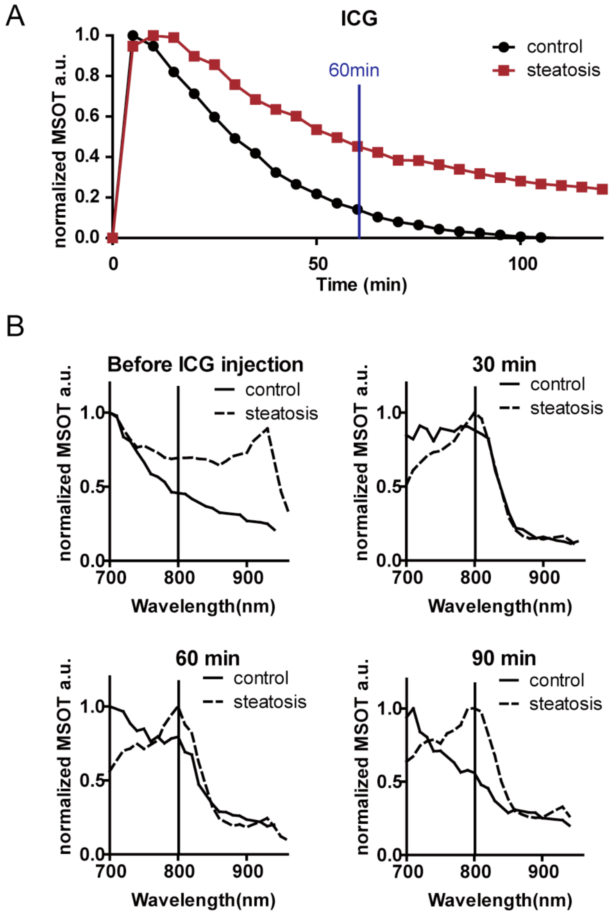


Fig.S2 Longitudinal monitoring of hepatic ICG clearance in mice. A. 2 hour monitoring of ICG intensity in control and steatotic livers (time interval: 5 minutes). n = 3 (3 positions in one animal). Time point data is normalized to the highest intensity acquired during

the observation. B Normalized spectra of control and steatotic liver before ICG injection, 30 min, 60 min and 90 min after ICG injection.

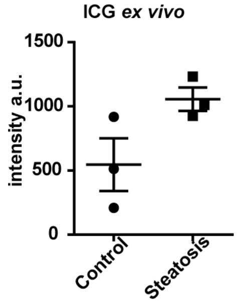


Fig.S3 *Ex vivo* quantification of ICG tracer. A. Quantification of mean residual ICG fluorescence intensity of the whole liver tissue section by fluorescence microscopy. n = 3.

Movie S1. Longitudinal ICG tracing in mouse livers by MSOT
The images are focus on lower abdominal region of mice. Time interval: 5 minutes. Total length: 120 minutes. Red: HbO2; blue: Hb; yellow: lipid; green: ICG.
